# The SIN3A histone deacetylase complex is required for a complete transcriptional response to hypoxia

**DOI:** 10.1101/182691

**Authors:** Maria Tiana, Barbara Acosta-Iborra, Laura Puente-Santamaría, Pablo Hernansanz-Agustin, Rebecca Worsley-Hunt, Norma Masson, Francisco García-Rio, David Mole, Peter Ratcliffe, Wyeth W. Wasserman, Benilde Jimenez, Luis del Peso

## Abstract

Cells adapt to environmental changes, including fluctuations in oxygen levels, through the induction of specific gene expression programs. To identify genes regulated by hypoxia at the transcriptional level, we pulse-labeled HUVEC cells with 4-thiouridine and sequenced nascent transcripts. Then, we searched genome-wide binding profiles from the ENCODE project for factors that correlated with changes in transcription and identified binding of several components of the Sin3A co-repressor complex, including SIN3A, SAP30 and HDAC1/2, proximal to genes repressed by hypoxia. *SIN3A* interference revealed that it participates in the downregulation of 75% of the hypoxia-repressed genes in endothelial cells. Unexpectedly, it also blunted the induction of 47% of the upregulated genes, suggesting a role for this corepressor in gene induction. In agreement, ChIP-seq experiments showed that SIN3A preferentially localizes to the promoter region of actively transcribed genes and that SIN3A signal was enriched in hypoxia-repressed genes, prior exposure to the stimulus. Importantly, SINA3 occupancy was not altered by hypoxia in spite of changes in H3K27ac signal. In summary, our results reveal a prominent role for SIN3A in the transcriptional response to hypoxia and suggest a model where modulation of the associated histone deacetylase activity, rather than its recruitment, determines the transcriptional output.

## INTRODUCTION

Organisms are constantly exposed to a wide range of environmental changes and maintenance of tissue homeostasis critically depends on transcriptional responses to different types of stresses. These responses typically involve the mobilization of hundreds of genes with roughly equal numbers of genes being up- and downregulated (1). Fluctuations in oxygen availability are a common stress that affects oxidative metabolism and the production of oxygen free radicals, thus metazoan cells constantly monitor oxygen levels. When oxygen metabolic requirement exceed its supply, a condition known as hypoxia, cells induce a transcriptional programme aimed at reducing oxygen consumption and increasing its delivery. This gene expression response is mainly driven by the Hypoxia Inducible Factors (HIFs), a family of heterodimeric transcription factors composed of alpha and beta subunits. The stability and transcriptional activity of HIF alpha subunit is regulated by oxygen, whereas HIF beta subunit (Ah Receptor Nuclear Translocator, ARNT), is insensitive to hypoxia. Of the three genes encoding for HIF alpha subunits, HIF1A (2) and EPAS1 (also known as HIF2A) (3–5) are the best characterized members of the family. The proteins encoded by these genes share a common mechanism of regulation by hypoxia that involves the control of stability and transactivation activity through oxygen-dependent hydroxylation of specific residues (6, 7). However, these genes differ in their pattern of expression. While *HIF1A* gene is ubiquitously expressed, *EPAS1* mRNA is restricted to specific tissues and cell types; being particularly abundant in endothelial cells (3, 4, 6). In addition, these factors regulate partially overlapping sets of target genes (8, 9). The HIF pathway is activated in a wide variety of pathological situations, including highly prevalent diseases such as cancer and cardiorespiratory diseases (10). Thus, understanding how hypoxia regulates gene expression could open up new avenues for clinical management of these diseases.

Several independent reports have investigated the regulation of gene expression in response to hypoxia from a global perspective. A consistent finding in these studies is that, although HIFs are required for both gene induction and repression (11), their DNA binding profile only correlates with gene induction; suggesting that repression is indirectly mediated by HIF (9, 12–18). In keeping with this hypothesis, HIF1A directly regulates the expression of repressors such as *MXI1* (19), *BHLHE40 (20)* and *BACH1* (21), that in turn repress the expression of individual genes in response to hypoxia (21–24). However, the relative contribution of these factors to hypoxia-induced repression is unknown and the general mechanism by which hypoxia represses transcription is not clear yet. On the other hand, although the mechanism of gene upregulation is much better understood, only a relatively small fraction (~10-20%) of the genes induced by hypoxia have a proximal HIF binding site (13, 25). Although these genes could be directly regulated by HIF bound to a distant enhancer, their regulation could also be indirectly mediated by transcription factors acting downstream of HIF. In agreement with this possibility, HIF induces the expression of many transcription factors (13, 26), although their role on the gene regulatory network induced by hypoxia is unclear yet. Thus, in spite of important advances, relevant questions regarding the mechanism by which hypoxia regulates transcription, and in particular gene repression, remain unanswered.

A further aspect of the transcriptional response to hypoxia where our knowledge is scarce regards to the role of cofactors (27). The role of p300/CBP as a HIF coactivator necessary for gene induction has long been stablished (28) and the interaction between these proteins is one of the oxygen-regulated steps in the transcriptional response to hypoxia (29). However, p300/CBP is only required for 35-50% of global HIF-1-responsive gene expression (30). Other cofactors, including CDK8 that is enriched at the regulatory regions of ~65% of hypoxia-inducible genes (31), are thus likely to be required for a complete response. The role of corepressor complexes in the response to hypoxia has not been addressed yet, although it is known that HIF recruits histone deacetylases (HDACs) and, intriguingly, HDAC inhibitors prevent HIF-mediated transcription (reviewed in (32)). Several corepressor complexes have been identified and it is generally assumed that gene repression is achieved by the combinatorial action of various enzymatic corepressor complexes that are recruited to DNA by sequence-specific transcription factors (repressors) that often act through enzymatic modification of histone protein tails (33). However, in spite of their role in transcription silencing, mounting evidence suggest that corepressors may have a much broader role in the regulation of gene expression including the induction of transcription (34). One of the best studied is the SIN3 complex, a conserved corepressor found from yeast to animals, that associates with histone deacetylases HDAC1 and HDAC2 (35, 36). Interestingly, SIN3A interacts with the repressors MXI1 (58) and E2F7 (59, 60) that act downstream of HIF to promote gene repression in response to hypoxia.

In this work we aimed to identify factors that contribute to control the gene expression program in response to hypoxia and centered our analysis in endothelium as a relevant cell type for the induction of angiogenesis, one of the key adaptive responses to hypoxia. Through the analysis of the binding profile of almost 200 factors generated by the ENCODE project, we found that several subunits of the SIN3A complex were overrepresented in the proximity of genes whose transcription is repressed by hypoxia. Further analysis demonstrated that SIN3A was required for full repression of 75-91% of genes downregulated by hypoxia. Unexpectedly, we also found that SIN3A was also required for the complete induction of 47-51% of upregulated genes. Thus, SIN3 regulates the vast majority of the transcriptional response to hypoxia. The analysis of the genome-wide binding profile of SIN3A shows that the complex is present in the promoter of hypoxia-regulated genes in normoxia and its binding to these loci is not altered by hypoxia, unlike H3K27ac signal that changes in parallel to the effects of hypoxia in gene expression. Therefore, in contrast to a simple model where repression is regulated by targeting of the corepressor complex, our analysis unveils that the recruitment of SIN3A is not required as a regulatory step in the transcriptional response to hypoxia. More generally, our results suggest that SIN3A is a general modulator of transcription that is essential to mount a complete response to environmental stresses through a mechanism that does not involve differential recruitment to target loci.

## MATERIAL AND METHODS

### Cell culture and treatments

Human Umbilical Vein Endothelial Cells (HUVEC) were purchased from Lonza (Lonza, C2519A) and maintained in Endothelial Cell Growth Medium (EGM-2 Bullet Kit, Lonza, CC-3162). HEK 293T were maintained in Dulbecco’s modified Eagle’s medium (Gibco, 41966052) supplemented with 50 U/ml penicillin (Gibco, 15140122), 50 μg/ml streptomycin (Gibco, 15140122), 2 mM glutamine (Gibco, 25030123) and 10% (v/v) fetal bovine serum (Gibco, 10270106). All cells were grown at 37°C and 5% CO2 in a humidified incubator containing 5% CO2 and tested regularly for mycoplasma contamination. For hypoxia treatment, cells were grown at 37°C in a 1% O2, 5% CO2, 94% N2 gas mixture in a Whitley Hypoxystation H35 (Don Withley Scientific).

### Metabolic labelling with 4-thiouridine and purification of newly synthesized mRNA

We used the protocol described by Dolken et al, and Schawnhausser et al (37, 38) that is represented in the Figure 1A. Briefly, exponentially growing HUVEC cells were exposed to 21% or 1% oxygen for 8 hours and pulse‐labelled with 4-thiouridine (400 μM, 4sU, Sigma, T4509) during the last two hours of treatment. After treatment, total cellular RNA was isolated from cells using TRI-reagent (Ambion, AM9738). 100 μg of total RNA was subjected to a biotinylation reaction to label the newly transcribed RNA containing the 4sU moiety (Pierce, EZ-Link Biotin-HPDP, 21341). Then, RNA was purified using Ultrapure TM Phenol:Chloroform:Isoamylalcohol (Invitrogen, 15593‐031) and labelled RNA was isolated from the total RNA by affinity chromatography using streptavidin coated magnetic beads (μMacs Streptavidin Kit; Miltenyi, 130‐074‐101).

**Figure 1.**
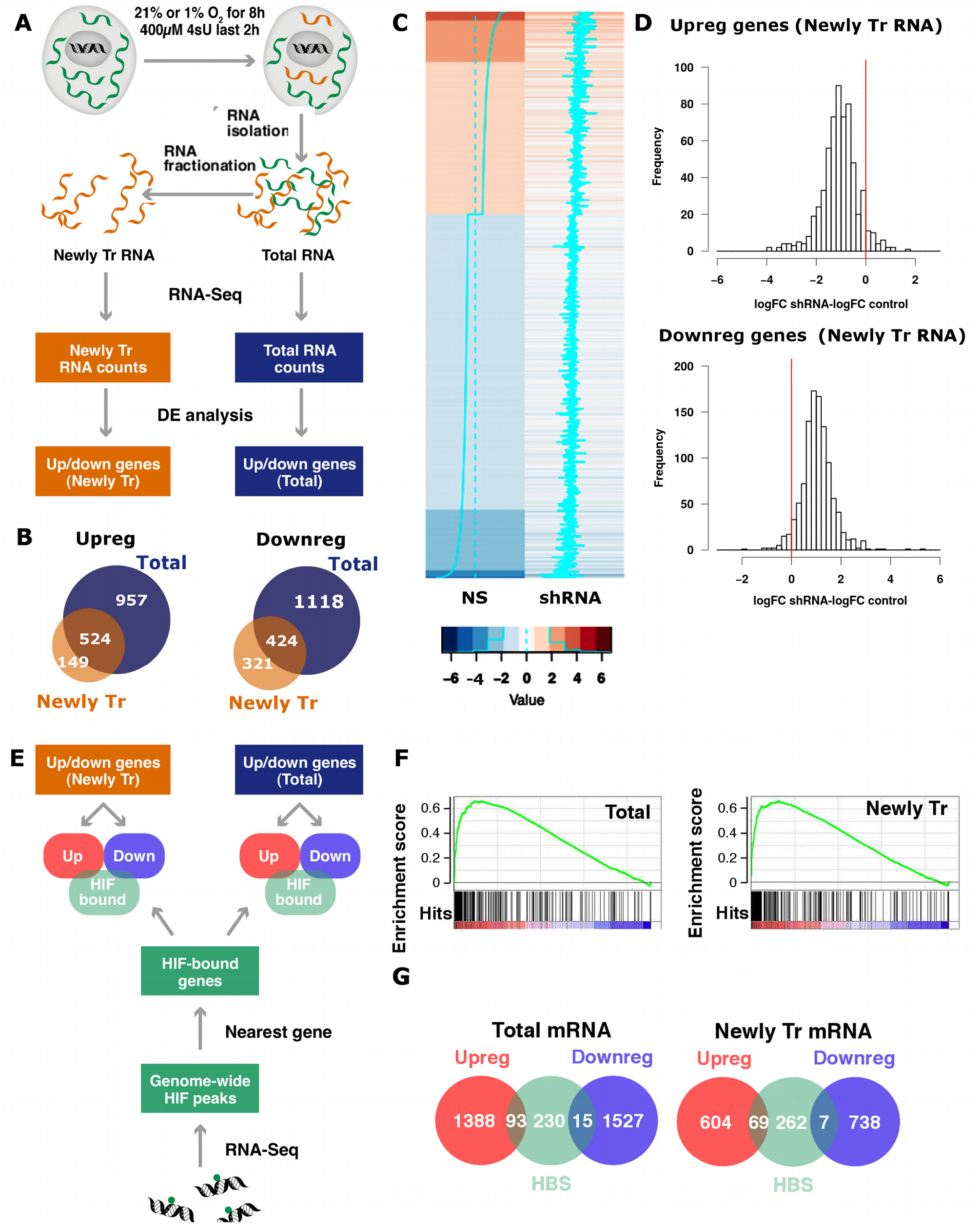
Identification of the transcriptional component of the hypoxia-induced gene expression pattern. **(A)** Exponentially growing non-synchronized HUVEC were exposed to 21% or 1% oxygen for 8 hours and pulse labelled with 4-thiouridine (4sU) prior RNA extraction. An aliquot of each RNA sample was kept (Total RNA) and the rest was processed to isolate the newly transcribed (4sU-labelled) RNA pool. PolyA RNA was purified from each of the three fractions (total, 4sU-labelled and pre-existing RNA), analyzed by high-throughput sequencing and quantified by mapping reads to the exonic regions of genes (RNAseq). The results shown derive from three independent biological replicas, each using a different pool of HUVEC donors. **(B)** Venn diagrams representing the number of genes significantly (FDR<0.05) up- or down-regulated shared by the total (blue circles) and newly transcribed (orange circles) fractions. **(C, D)** Control (NS) and *EPAS1*-silenced (shRNA) HUVEC were exposed to 1% oxygen for 16 hours, pulse labelled with 4-thiouridine (4sU) and the newly transcribed mRNA fraction from each sample was analyzed by RNAseq. The results shown derive from a single biological replica. The treatment with shRNA reduced the expression of *EPAS1* to 17% of control values. **(C)** Heatmap representing the log-fold change (logFC) expression in hypoxia over normoxia in control (“NS”) and *EPAS1* knockdown cells. (“shRNA”) for all transcripts affected by hypoxia (absolute logFC>1). Each row represents a transcript in a colour code that indicates their response to hypoxia (upregulated genes in red and downregulated in blue) and the magnitude of change as shown indicated in the legend below the heatmaps. The cyan lines also indicate the magnitude of the fold change for a particular transcript, and the dotted line indicates no change in expression. **(D)** For each gene whose expression up-(logFC>1, top panel) or down-regulated (logFC< −1, bottom panel) by hypoxia in control cells, we computed the difference between the changes induced by hypoxia in *EPAS1*-silenced cells and control cells, that is the logFC observed in *EPAS1*-silenced cells minus logFC observed in controls (“logFC shRNA-logFC NS”). The red line indicates a difference value of zero, which would be expected in the case of lack of effect of the interference. The mean of the differences was significantly different of zero in all the cases (single sample student’s t-test, p<0.001). **(E)** HUVEC were exposed to 1% oxygen for 16 hours and then processed to determine ARNT and EPAS1 binding sites by ChIP-Seq. Regions bound by HIF were assigned to the nearest gene and compared to mRNAs significantly up-(“Upreg.”) or down-regulated (“Downreg.”) by hypoxia in the total and 4sU-labelled fractions from part A. **(F)** The genes detected by RNA-seq in total (left graph) and newly transcribed fractions (right graph) were sorted according to their response to hypoxia (from strongly induced on the left, labelled in red colour, to strongly repressed on the right, labelled in blue colour) and distribution of EPAS1 binding sites across these ranked lists of genes was analysed by Gene Set Enrichment Analysis (GSEA). HIF binding sites were significantly enriched (FDR<0.01) in the genes whose expression was induced by hypoxia in both mRNA fractions. (G) Venn diagrams representing the number of genes significantly (FDR<0.05) up-(red circles) or down-regulated (blue circles) by hypoxia that present a HIF binding site (HBS, green circles). The same analysis was performed for genes differentially expressed in the total mRNA (left graph) and 4sU-labelled (right graph) fractions. The proportion of genes showing HIF binding sites was significantly different for the up- and downregulated categories (two-proportion test, p<0.001).

### High-throughput sequencing and bioinformatics analysis

Libraries were prepared using the standard protocol for messenger RNA sequencing (Illumina, TruSeq Stranded mRNA) and sequenced on HiSeq2000 instrument (Illumina) according to the manufacturer’s protocol. The resulting reads were aligned to the GRCh37/hg19 assembly of the human genome using TopHat (39) with the default parameters. Finally, the gene expression level was calculated as the number of reads per gene, computed using HTSeq (40) and gene features as defined in the GRCh37.75 release of the human genome (gtf file) and expressed as RPKMs. Differential gene expression analysis was performed with the Bioconductor (41) edgeR package (42) for the R statistical Software (http://www.R-project.org/). The raw reads and processed data derived from RNA‐seq experiments are available at NCBI’s Gene Expression Omnibus (43, 44) and are accessible through the following GEO Series accession numbers: GSE89831, GSE89838, GSE89839, GSE89840. All computations were performed using R software package (http://www.R-project.org/)

### Chromatin immunoprecipitation

Chromatin immunoprecipitation was performed as previously described by Schödel et al (14) using antibodies against HIF1A (PM14), EPAS1 (PM9) (9, 45), ARNT (Novus, NB-100-110; RRID:AB_10003150), SIN3A (Abcam, ab3479; RRID:AB_303839) and a Rabbit Control IgG (Abcam, ab46540; RRID:AB_2614925).

Libraries were prepared using the Illumina ChIP-seq kit and sequenced on the Illumina GAII platform according to the manufacturer’s protocol. Sequences were mapped to the GRCh37/hg19 assembly of the human genome using ELAND software (HIF ChIPseq) or Bowtie2 (REF) (SIN3A ChIPseq). Binding peaks were determined from the aligned reads using MACS software (46) using default parameters and the preimmune sample as a control. A single ChIP experiment was analysed by ChIP-sequencing and Raw and mapped data are available at GEO (GSE89836 and GSE103245). The analysis of mapped reads and genomic intervals defined by the bound regions was performed with the GenomicRanges (47) and Genomation (48) Bioconductor packages.

### Lentiviral Production

Lentiviral vector pGIPz (Open Biosystems) was used to silence the mRNA expression of HIF1A (V2LHS_132150), EPAS1 (V2LHS_113753), SIN3A (V3LHS_343545) and MXI1 (V3LHS_408351), as well as a Non-silencing lentiviral shRNA Control (NS; RHS4346). Lentiviruses were produced and tittered in HUVEC cells as previously described (49). For transduction of target cells, lentiviruses were used at a multiplicity of infection (MOI) of 1-2 for 8h, resulting in more than 95% transduced (GFP-positive) cells 72 hours after infection.

### Identification of trans-acting factors enriched in transcriptionally regulated genes

The March 2012 internal data freeze consisting in 690 ChIP-seq datasets representing 161 unique regulatory factors and generated by the ENCODE Consortium (50) was downloaded from the UCSC genome browser (51, 52) and from Bioconductor using AnnotationHub version 2.4.2. We only considered peaks with −log10 FDR>2. For transcription factors that had been tested in more than one cell line only the intersection of all experiments was considered. HIF1A binding sites conserved across cell types obtained from the integration of published ChIP-chip and ChIP-seq experiments data (12, 13, 15, 18, 53) and HUVEC genome-wide EPAS1 binding sites derived from our ChIP-seq were also included. Peaks of transcription factors binding were ascribed to the closest transcription start site. Significantly enriched binding sites for any of the transcription factors were determined by a Fisher’s exact test. All analyses were performed in R software package (http://www.R-project.org/).

## RESULTS

### Transcriptional regulation in hypoxia is indirectly mediated by HIF

HIFs are master regulators of the gene expression pattern induced by hypoxia that are required for the induction and repression of the vast majority of genes regulated by hypoxia (11). However, many of the effects of HIF on gene expression could be indirect since only ~10-20% of the induced genes present HIF binding sites (13, 25) and HIF binding does not correlate with gene repression (13, 14, 17, 25). Therefore, we sought to identify transcription factors acting downstream of HIF during the adaptive response to hypoxia of Human Umbilical Vein Endothelial Cells (HUVEC).

The gene profiling studies published to date were based on the analysis of steady state mRNA levels, and thus cannot differentiate between effects on transcription and decay rates. To address this caveat, we pulse-treated HUVEC for two hours with 4-thiouridine (4sU) to label nascent RNA and then characterized the pattern of newly transcribed mRNAs by affinity capture of the labelled transcripts followed by high-throughput sequencing (figure 1A, top). Incorporation of 4sU into mRNA after a short pulse can be used as a proxy for transcription rate (37, 38) and comparison of changes in this nascent fraction to those in total RNA allowed us to identify and focus on those genes whose regulation by hypoxia occurs at the transcriptional level (figure 1A). This analysis identified 1481 and 673 upregulated genes (FDR <0.05) in the total and newly transcribed mRNA fractions respectively and 1542 and 745 downregulated genes in these same fractions (figure 1B and supplementary table 1). Comparison of the genes differentially expressed in each of these fractions revealed that only 35.4% of the genes whose expression was induced by hypoxia showed statistically significant changes in their transcription rate. Likewise, only 27.5% of downregulated genes had significantly repressed their transcription. Thus, although transcription is the major determinant of the gene expression changes induced by hypoxia (Tiana et al. manuscript in preparation), we only found statistically significant changes in transcription rate for a subset of the genes showing altered steady state expression levels.

We next decided to investigate the role of HIF on the changes in transcription induced by hypoxia and focussed on EPAS1 which is the predominant HIF isoform expressed in HUVEC (supplementary figure 1). Figure 1C represents the effect of hypoxia on transcription (logarithm of the ratio of nascent mRNA levels in hypoxia over normoxia) in control (“NS”) and *EPAS1* silenced cells (“shEPAS1”, knock-down to 15-19.8% of control values) as a heat map where each row represents an individual gene and the color code indicates the magnitude and direction of the changes induced by hypoxia (log-fold change hypoxia over normoxia). To quantify the magnitude of the effect, we first classified genes as up-regulated (log-fold change>1) or downregulated (log-fold change< −1) attending to the effect of hypoxia in their transcription rate in control cells. Then, for each gene, we calculated the difference of log-fold induction values in EPAS1-silenced (figure 1D, “shRNA”) minus control cells (Figure 1D, “NS”). Of the 617 genes whose transcription was induced by hypoxia in control HUVEC, 581 (94%) showed reduced expression in knock-down cells (Figure 1D top panel; mean of differences= −1.05, 95% CI [−1.11, −0.99]). Similarly, *EPAS1*-silencing attenuated the transcriptional repression of 1021 out of 1067 genes (96%), resulting in positive values of fold induction differences (Figure 1D bottom panel; mean of differences= +1.04, 95% CI [+1.01, +1.08]).

These experiments indicate that EPAS1 globally mediates the effects of hypoxia on transcription in HUVEC, affecting both induction and repression to a similar extent. Next, to investigate if the effects on transcription were directly mediated by HIF we determined the genome-wide binding patterns of EPAS1 and ARNT in HUVEC by chromatin immunoprecipitation followed by high-throughput sequencing (ChIP-seq) (figure 1E). To increase the stringency in the annotation we only considered genomic segments bound by both EPAS1 and ARNT, as proposed in previous studies (14). This analysis yielded a total of 392 regions bound by ARNT/EPAS1 in HUVEC (supplementary table 2), a value of similar magnitude to that found in other cell types (12, 13, 17). In order to compare pan-genomic binding data with the effects of hypoxia on gene expression, EPAS1 binding regions were assigned to the nearest gene locus, resulting in a total of 338 genes bound by this factor (supplementary table 2). EPAS1 bound genes strongly clustered with genes up-regulated by hypoxia in a Gene Set Enrichment Analysis (GSEA) (54) in both total (figure 1F, left graph) and newly transcribed (figure 1F, right graph) mRNA fractions; strongly supporting that HIF binding increases transcription rate. However, only 6% (93 out of 1481) and 10% (69 out of 673) of loci bound by EPAS1 were significantly induced by hypoxia in the total and newly transcribed fractions respectively (figure 1G). In the case of gene repression, 1% of the genes with reduced expression in total (15 out of 1527) or newly-transcribed (7 out of 745) fractions were bound by EPAS1 (figure 1G). This proportion is significantly lower than the 6%-10% observed for upregulated genes (normal approximation of proportions, Student’s t-test, p<0.01). Thus, almost all the transcriptional repression and, at least part, of the induction are indirectly mediated by HIF in HUVEC.

### Identification of candidate DNA-binding proteins contributing to gene regulation in hypoxia

In order to identify the DNA-binding proteins that could mediate the indirect effects of HIF on transcription, we employed motif finding algorithms (55–57) to search for factors over-represented in the vicinity of genes whose transcription was significantly affected by hypoxia. However, this approach is limited by design to sequence-specific transcription factors (TF) and shows low specificity (58).

To overcome these limitations and expand our search to coregulatory proteins binding DNA indirectly, we exploited the hundreds of ChIP-seq experiments from the ENCODE Consortium (50). Specifically, we searched for trans-acting factors over-represented in the regulatory regions of genes whose transcription is repressed by hypoxia (figure 2A). Since the number of factors experimentally determined in HUVEC is small, we used binding sites of all transcription factors analysed by ENCODE in any cell line as a proxy for the genome-wide regulatory landscape. To exclude cell-type specific binding events and maximize the selection of sites likely to be present in HUVEC, for each TF we only considered binding sites that were conserved (i.e. overlapping) in all cell lines studied by the ENCODE consortium (figure 2A). These stringent criteria resulted in a list of binding sites expected to be conserved across cell lines. In addition to the factors analysed by ENCODE, we also included a data set describing HIF1A, EPAS1 and ARNT binding sites derived from the integration of published ChIP-chip and ChIP-seq experiments (59). Finally, we also included into this analysis pipeline the genome-wide EPAS1 binding sites derived from the ChIP-seq experiment described herein. For each DNA-interacting factor, we ascribed binding sites to the nearest transcription start site and compared the distribution of binding sites in genes induced or repressed by hypoxia with those whose transcription remained constant (figure 2A).

**Figure 2.**
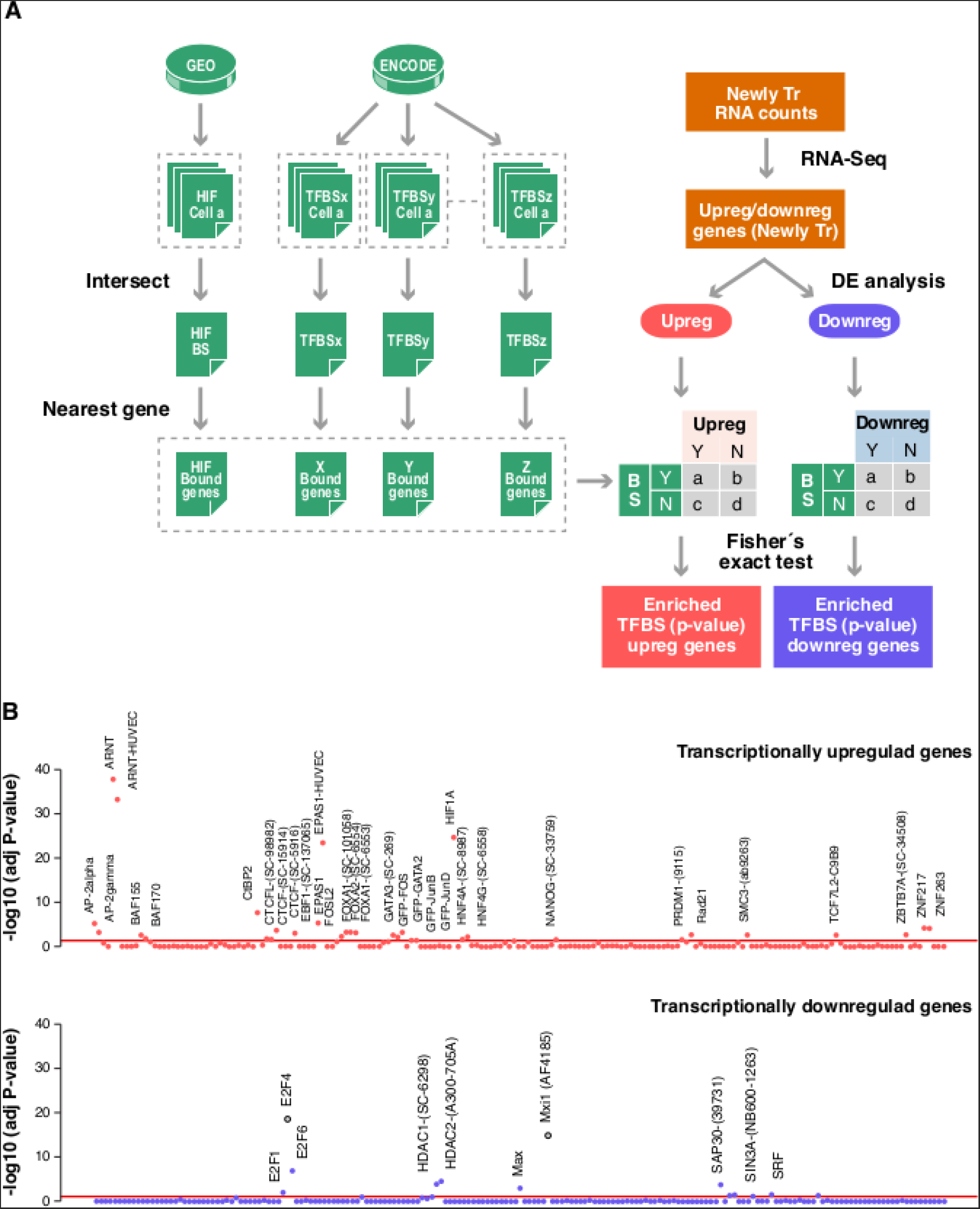
Transcription factor binding sites over-represented in genes regulated by hypoxia. The presence of binding sites for each one of the 166 factors (“TFBSx”, “TFBSy”,…“TFBSz”) assayed by the ENCODE project and HIF binding sites determined elsewhere (“HIF”) as determined for each of the genes whose transcription is significantly modulated by hypoxia in HUVEC cells (“Newly Tr. RNA counts”). For factors assayed in more than one cell line by ENCODE, we only considered those peaks present in all cell types. For each transcription factor we assigned binding sites to the nearest gene. For association studies, genes were categorized according to their response to hypoxia into up-or down-regulated sets and the distribution of binding sites for each factor in each set was compared to that expected by chance by Fisher exact test. The resulting p-values were adjusted for multiple testing. **(A)** Schematic diagram depicting the analysis strategy. **(B)** The figure shows the association between binding for each of the analysed factors (x-axis) and the induction (upper graph) or repression (bottom graph) of the nearest genes. Each dot represents the FDR-adjusted p-value obtained in the Fisher’s for a single factor. Red line correspond to an adjusted p-value of 0.005. The identity of the factors significantly over-represented is indicated.

As expected, this analysis revealed that the distribution of HIF1A, EPAS1 and ARNT binding sites was strongly biased toward genes whose transcription was induced by hypoxia (figure 2B upper graph, “transcriptionally upregulated genes”; Fisher’s exact test, FDR-adjusted p-value<0.0001). We also found enrichment for other factors, albeit showing a much less skewed distribution towards upregulated genes (figure 2B upper graph). Thus, these additional transcription factors could contribute to gene induction in hypoxia acting cooperatively with HIF or downstream of it.

Importantly, the analysis of genes whose transcription is repressed under hypoxia also revealed a set of DNA-binding factors that were significantly overrepresented (figure 2B bottom graph, “transcriptionally downregulated genes”). Among them was the transcriptional repressor MXI1, a factor known to act downstream of HIF to repress specific genes (19), the MXI binding partner MAX, and members of the E2F family, particularly the transcriptional repressors E2F4 and E2F6. In addition to these sequence-specific transcription factors, we also found significant enrichment for histone deacetylases 1 and 2 (HDAC1 and HDAC2) and the proteins SIN3 Transcription Regulator Family Member A (SIN3A) and Sin3A Associated Protein 30kDa (SAP30). Notably, HDAC1/2, SAP30 and SIN3A are components of the histone deacetylase co-repressor complex Sin3A which is required for transcriptional repression by MXI1 (60) and E2Fs (61, 62). Taken together, these results suggest that the Sin3A co-repressor complex could mediate the repression of transcription observed under hypoxia.

### Depletion of SIN3A disrupts the hypoxic gene expression pattern

In the view of these results, we decided to focus our attention on the role of the SIN3A complex on the transcriptional response to hypoxia. To investigate the functional role of this complex, we analysed the effect of *SIN3A* knock-down on the hypoxic transcriptome by means of RNA-sequencing (figure 3 and supplementary table 3). Figure 3A represents the effect of hypoxia on transcription in control (“NS”) and *SIN3A* silenced cells (“shRNA”) as a heat map where each row represents an individual gene and the color code indicates the magnitude and direction of the changes induced by hypoxia (log-fold change hypoxia over normoxia). *SIN3A* knock-down to 36% of control levels had a profound impact on the transcriptional response to hypoxia and, unexpectedly, it affected not only gene repression but also impaired gene induction. To quantify the magnitude of the effect, we first classified genes as upregulated (FDR<0.001 and log-fold change>0) or downregulated (FDR<0.001 and log-fold change< 0) attending to the effect of hypoxia in the mRNA levels in control cells. Then, for each gene, we calculated the difference of log-fold induction values (log2 hypoxic over normoxic mRNA values) in *SIN3A*-silenced minus their value in control cells. *SIN3A* RNA interference attenuated the repressive effect of hypoxia for 779 out of the 861 genes (91%) that were downregulated in control cells (figure 3B, bottom graph; mean of differences= +0.41, 95% CI [+0.38,+0.43]). In agreement, less than 25% (211 out of 861) of the genes significantly repressed by hypoxia in control conditions were still found significantly repressed in knockdown cells (figure 4C, bottom graph).Gene induction was also affected by *SIN3A* interference, with 661 out of 816 upregulated genes showing reduced induction in knock-down cells (figure 3B, upper graph). This effect was of smaller magnitude compared to the effect on repressed genes (mean of differences= −0.27, 95% CI [−0.29,−0.25]), but statistically significant (single sample Student’s t-test, p<0.001). Accordingly, 53% (430 out of 816) of the genes induced in control cells were still significantly upregulated upon SIN3A interference (figure 3C, upper graph). In agreement with these results, analysis of the ontology terms associated to the genes regulated by hypoxia revealed that *SIN3A* is required for a wide range of biological pathways regulated by hypoxia (figure 3D). In particular, SIN3A interference affected metabolic functions repressed by hypoxia such as oxidative phosphorylation and fatty acid metabolism (figure 3D).

**Figure 3.**
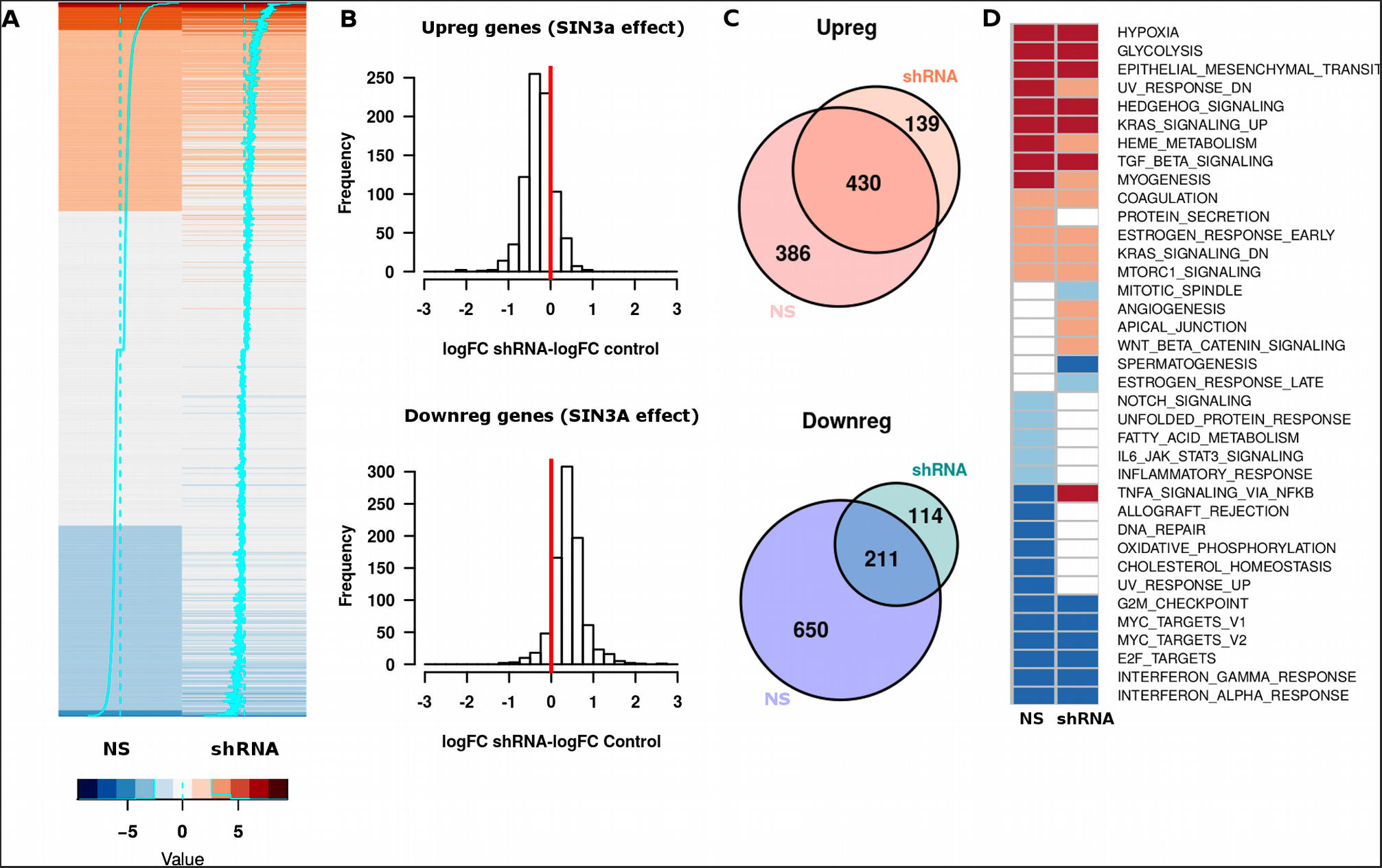
Effect of *SIN3A* knock-down on hypoxia-induced gene regulation. Exponentially growing non-synchronized HUVEC were transduced with lentiviral particles encoding for shRNA against *SIN3A* mRNA (“shRNA”) or for a non-specific shRNA (“NS”); 72 hours after infection cells were exposed to 21% or 1% oxygen for 16 hours, poly-A RNA was purified and analyzed by RNAseq. The results from two independent biological replicates, each using a different pool of HUVEC donors, were used to compute gene expression values. The treatment with shRNA reduced the expression of *SIN3A* to 34% (replicate 1) and 38% (replicate 2) of control values. The effect of hypoxia on each gene was calculated as the log-fold change of hypoxic over normoxic values (“logFC”). **(A)** Heatmap representing the changes in expression, as log2 fold changes of hypoxic compared with normoxic conditions (logFC), in control (“NS”) and SIN3A-silenced cells (“shRNA”). The graph represents all transcripts differentially expressed in control cells (FDR<0.01) as was described in figure 2. **(B)** For each gene whose expression was significantly (FDR<0.01) up-(upper graph) or down-regulated (bottom graph) by hypoxia in control cells, we computed the difference between the changes induced by hypoxia in *SIN3A*-silenced cells and control cells (“logFC shRNA-logFC NS”). The red line indicates a difference value of zero, which would be expected in the case of lack of effect of the interference. The mean of the differences was significantly different of zero in all the cases (single sample student’s t-test, p<0.001). **(C)** Venn diagrams representing the number of genes significantly up-(upper graph) or downregulated (bottom graph) in control and *SIN3A* (“shRNA”) interfered cells. **(D)** The heatmap represents the biological functions associated to genes differentially induced (represented in red) and repressed (blue) in control (“NS”) and silenced cells (“ShRNA”) as determined by GSEA against the “Hallmarks” gene sets. Light colors represent adjusted p-value<0.05 and dark colors adjusted p-value<0.01.

### SIN3A binding profile is not altered by hypoxia

The effect of SIN3A on gene induction was unanticipated given its established function as a corepressor and the observed association with genes repressed under hypoxia (figure 2B). In addition, hypoxia did not alter the expression of *SIN3A* nor affected its subcellular localization (supplementary figure 2). For these reasons, we next determined the genome-wide binding pattern of SIN3A in HUVEC. Sequencing of the chromatin immunoprecipitated with SIN3A revealed 16875 regions (FDR<5%) bound by this factor in normoxic HUVEC (supplementary table 4A), which is similar to the number of sites described by the ENCODE project in other cell lines (mean 11617 peaks, range 6024-21309 peaks, n=9 datasets). Interestingly, the analysis of the distribution of these regions across the different functional elements defined by the genome-wide segmentation of HUVEC (46), showed a strong enrichment of SIN3A in active promoter regions (figure 4A), with a distribution that was significantly different to that expected by chance (Chi-square_6_=213740, p<10^−15^). The correlation between SIN3A signal and gene expression, is further analyzed in figure 4B, which represents the SIN3A signal in a region of 4Kb centered at the transcription start site (TSS) across all human genes (each row represents a gene) sorted according to their expression in HUVEC. This representation shows that SIN3A signal is prominent on the promoters of genes actively transcribed in normoxic HUVEC but faint on genes with very low (1-10 counts per million of reads) or not detectable expression (less than 1 count per million of reads) (figure 4B). The same pattern was found when we analyzed the 11948 regions (FDR<5%) bound by SIN3A in hypoxic conditions (figure 4B and supplementary table 4B). Although paradoxical, this result concurs with previous reports describing the presence of SIN3A in active promoters (63) and the fact that HDACs are highly recruited to actively transcribed genes (64). Next, we studied whether the distribution of SIN3A correlated with the changes on transcription induced by hypoxia. To this end we selected genes expressed in HUVEC (those with cpm>1 in at least two samples of the RNA-seq experiments) and represented SIN3A signal across genes sorted by their response to hypoxia (log-fold change of the ratio hypoxia over normoxia) (figure 4C). Although we detected SIN3A in the promoters of the majority of expressed genes, the distribution of SIN3A binding sites was slightly skewed towards genes repressed by hypoxia (figure 4C). The bias in distribution of SIN3A signal with respect to hypoxia response is more clearly observed when SIN3A binding loci were examined for enrichment among hypoxia-regulated genes using GSEA (figure 4D and supplementary table 5). Interestingly, the skewed distribution was observed in normoxic cells prior to the exposure to the stimuli (figure 4C left panel and blue line in figure 4D), in agreement with the enrichment of SIN3A proximal to genes repressed by hypoxia found in the analysis of the ENCODE datasets (figures 2B and 4D). Moreover, the binding pattern of SIN3A under hypoxia (figure 4C, right panel, and red line in figure 4D) was very similar to the one observed under normoxic conditions. In agreement, we did not observe statistically significant differences in the recruitment of SIN3A to hypoxia-responsive genes (figure 4E). These results strongly argue against the regulation of SIN3A recruitment and point to modulation of the activity of the pre-assembled complex in response to hypoxia. Consistent with this possibility, the transcription of yeast genes *INO1* and *CHO* in response to nutrients was found to be mediated by modulation of the associated HDAC activity rather than recruitment of the SIN3A complex (63). To test this possibility, we analyzed publicly available datasets (GSE50144) of H3K27 acetylation in HUVEC (65) and found that, in response to hypoxia, the signal of this histone modification was increased in the promoters of induced genes and reduced in repressed genes (figure 4F). Altogether these results suggest that the activity of the complex, rather than its recruitment to target genes, could be regulated in response to hypoxia to modulate the transcription of the hypoxia-responsive genes.

**Figure 4.**
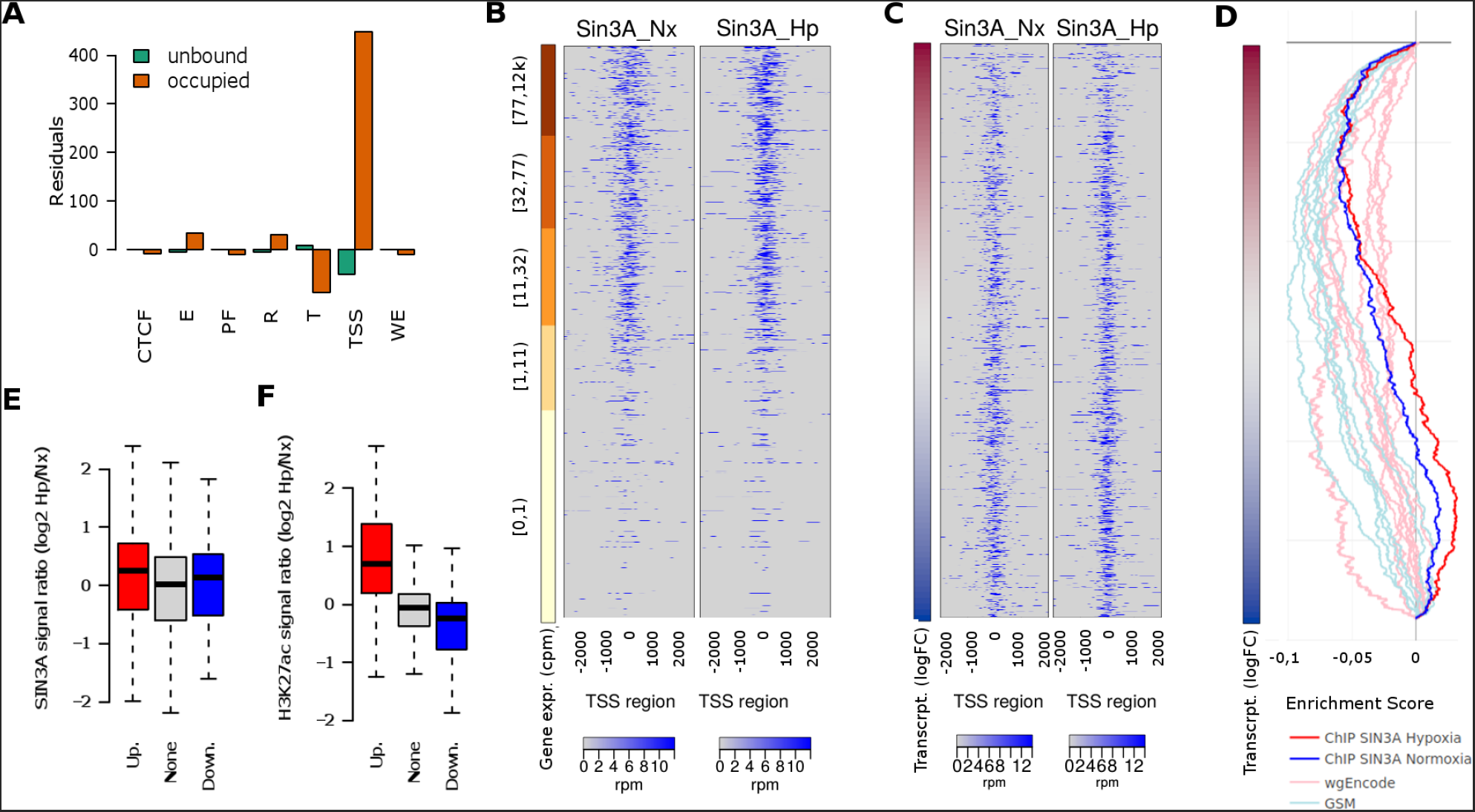
Genome-wide SIN3A binding profile. Exponentially growing non-synchronized HUVEC were exposed to 1% oxygen for 16 hours and then processed to determine SIN3A binding profile by ChIP-Seq. **(A)** The distribution of the normoxic SIN3A binding regions across the functional regions of the HUVEC genome was studied and compared to that expected by chance using a chi-squared goodness of fit test (Chi-square_6_=213740, p<10^−15^). The graph represents the Pearson residuals resulting from the test. The functional regions (chromatin states as defined by the ENCODE consortium) are: “TSS”, predicted promoter region including TSS; “PF”, Predicted promoter flanking region; “E”, Predicted enhancer; “WE”, Predicted weak enhancer or open chromatin cis regulatory element; “CTCF”, CTCF enriched element; “T”, Predicted transcribed region; “R”, Predicted Repressed or Low Activity region. **(B, C)** Heatmaps showing the enrichment of SIN3A in a 4 Kb region centered in the TSS. Each row represent a gene and rows are sorted according to gene expression level as determined from RNA-seq (B) or log fold change in transcription (Newly transcribed fraction) induced by hypoxia (C). In the latter case only genes with detectable expression (cpm>1 in at least two samples) are represented. **(D)** GSEA graph the distribution of the normoxic SIN3A (blue) and hypoxic (red) SIN3A binding sites from this study (ChIP SIN3A) and those determined by the ENCODE project (“wgEncode”, ligth red) and individual research groups (“GSM”, light blue) across genes expressed in HUVEC sorted according to their transcriptional response to hypoxia as in C. The information about each SIN3A data set together with ES and p-values for echrichment is shown in supplementary table 4. **(E)** The log-ratio of the normalized SIN3A signal in hypoxia over normoxia was computed for all promoters. The boxplot represent the distribution of values obtained for genes upregulated (“Up.”, logFC > 1 and adjusted p-values<0.001), downregulated (“Down.”, logFC < −1 and adjusted p-values<0.001) and not effected by hypoxia (“None”). The differences between groups were not statistically significant for alpha=0.01 (ANOVA F_2,9053_=3.707, p=0.0246). **(F)** Wig files from reference (53) were downloaded from the NCBI GEO repository (GSE50144). The log-ratio of the normalized H3K27ac signal in hypoxia over normoxia was computed for all promoters and represented as in (D). The differences between mean signals in each group were statistically significant (ANOVA F_2,9053_=247.9, p<2e-16) and the signal in both up- and down-regulated groups was significantly different to that in control genes (post-hoc Tukey HDS, adjusted p-value<2e-16).

## DISCUSSION

In this work we aimed to provide novel insights into the transcriptional response to hypoxia, a type of stress commonly found in a wide variety of physiological and pathological situations. Specifically, our objective was to identify novel transcription factors involved in this response.

To this end we employed a technique that, in contrast with previous studies, allowed us to determine the effect of hypoxia on transcription, avoiding the confounding effect that RNA decay rates might have on the gene expression pattern. This experimental approach identified a set of genes whose expression was regulated by hypoxia at the level of transcription and thus optimal for the identification of transcription factors that could cooperate with HIF or mediate its effects. Then, we exploited datasets of genome-wide binding profiles generated by ENCODE, to search for factors enriched in the proximity of hypoxia-regulated targets. Although ENCODE experiments were performed under normoxic conditions, we hypothesized that any hypoxia-regulated factor is likely to be present to some extent under normoxia and hence to be detected on its target genes. The comparison of the fraction of hypoxia-regulated and control genes bound by each factor led us to the identification of sets of proteins significantly enriched in genes transcriptionally induced or repressed by hypoxia. In addition to HIF1A, EPAS1 and ARNT, we identified 24 additional factors overrepresented in upregulated genes. Among them, we found FOS and its binding partners JUND and JUNB, in agreement with previous reports suggesting a cooperation between HIF and AP-1 in the induction of hypoxia-regulated genes (9, 53, 66). In the case of downregulated genes, we found enrichment for ten proteins, including several repressors of the E2F family, MXI1 and its dimerization partner MAX, as well as several components of the SIN3A histone deacetylase corepressor complex. These findings are particularly relevant as the mechanisms that mediate gene repression by hypoxia are poorly understood and the role of corepressors in the transcriptional response to hypoxia has not been studied before. For these reasons we decided to further investigate the role of the SIN3A corepressor.

Knock-down of *SIN3A* had a profound impact on the transcriptional response to hypoxia, affecting to 75% of the genes downregulated by hypoxia and 47% of upregulated genes. However, it is important to note that in most cases *SIN3A* depletion did not fully prevent, but rather attenuated the effect of hypoxia on gene expression. This could be a consequence of incomplete *SIN3A* interference as we could only reduce its expression to 36% of control levels on average. However, it is well known that removal of corepressor function results in gene expression changes that tend to be subtle, suggesting that they function to fine-tune transcript levels rather than turn transcription on and off (34). Given the role of SIN3A as a corepressor, the effect of its interference on hypoxia-triggered gene induction was puzzling. In some cases, the attenuated induction was a consequence of increased basal gene expression upon *SIN3A* interference, providing a potential explanation for this effect that is in agreement with the function of SIN3A as a corepressor. However, this was not the case for the majority of genes (360 out of 386) showing attenuated induction, suggesting that SIN3A may be required for full induction of some genes. Although these results might seem paradoxical, SIN3A was initially described in yeast as a protein with dual functions as activator and repressor (61, 62) and recent works are putting forward its role as an activator of specific genes (reviewed in (29)). Of particular relevance to our work, previous works suggest that the SIN3A histone deacetylase complex is required for the induction of genes in response to environmental insults such as heat or osmotic stress (63, 64) in yeast and xenobiotics in mammalian cells (67). Thus, additionally to its undisputed role as corepressor, SIN3A seems to be also playing a role on transcriptional activation. In fact, the role of corepressors in transcriptional upregulation is not unique to SIN3A. Loss of different types of corepressor proteins usually results in similar numbers of genes decreasing and increasing their expression (34), so that emerging models propose that recruitment of both corepressor and coactivator complexes is needed for gene induction (33). Importantly, the requirement of SIN3A for hypoxia-triggered gene induction could explain, at least in part, why histone deacetylase inhibitors, that in most cases promote gene expression, in the context of several HIF target promoters prevent transcription (30, 68).

Aiming to understand the molecular mechanism by which SIN3A affects the transcriptional response to hypoxia we determined its genome-wide binding profile. Interestingly, we found that SIN3A is preferentially found at the promoter region of actively transcribed genes, in agreement with a recent report showing that SIN3A and NANOG co-occupy active promoters (69). Although, SIN3A signal was found in most active promoters, it was absent from about 30% of the hypoxia-inducible genes. In contrast, it was absent from 5% of repressed genes only; resulting in a distribution of SIN3A binding sites slightly skewed toward hypoxia-repressed genes. This distribution was highly reproducible as it was also observed in other SIN3A binding datasets, including those from ENCODE and individual GEO submissions (figure 4D and supplementary table 5). A question arising from these results is whether this pattern is shared by other corepressor complexes. In addition to SIN3A, ENCODE also determined the binding profiles of COREST, CTBP2, MTA3, SMARCA4, SMARCB1, SMARCC1, SMARCC2, MBD4, CHD1, CHD2, KDM5A, KDM5B and HDAC6 that are subunits of different corepressor and chromatin remodelling complexes such as CoREST, NURD and SWI/SNF. However, none of these factors were enriched in genes downregulated by hypoxia (figure 2). On the other hand, analysis of datasets generated by individual groups and deposited in GEO confirmed an over-representation of binding of components of the SIN3A complex but not subunits of the NCor, SMRT, NURD or CoREST corepressors in repressed genes (supplementary tables 5 and 6). Thus, SIN3A enrichment on the proximity of hypoxia repressed genes seems to be rather specific. Interestingly, the comparison of normoxic and hypoxic SIN3A binding profiles revealed very similar patterns suggesting that, in spite of the effect of *SIN3A* interference on gene expression, hypoxia did not regulate SIN3A recruitment. In agreement, although we found individual cases where SIN3A is recruited by hypoxia to repressed genes (supplementary figure 3), as a general rule we did not find significant changes in the SIN3A signal associated to promoters of upregulated nor downregulated genes (figure 4). In contrast, H3K27ac signal on the promoter of repressed genes was significantly reduced in response to hypoxia. Although we can not rule alternative mechanisms, our data support a model where hypoxia regulates the activity of SIN3A complexes constitutively located at target genes, rather that its recruitment. Supporting this possibility, a recent work demonstrates that yeast Sin3 can be detected at target promoters *INO1* and *CHO2* under repressing and derepressing conditions and contributes to the induction or repression of these genes through its interaction with activators or repressors that are recruited to these promoters under different conditions (63). Thus, a plausible explanation is that HIFs induce the expression of sequence-specific repressors that in turn activate the histone deacetylase activity of the SIN3A complex already bound to target genes. In this regard, the SIN3A complex interacts to MXI1 to silence gene expression (70) and we found enrichment of MXI1 binding to downregulated genes (figure 3). Moreover, *MXI1* mRNA is robustly induced by hypoxia in HUVEC cells (supplementary table 1) in a HIF-dependent manner (data not shown). To investigate the possibility of MXI1 acting downstream of HIF to regulate SIN3A located at target genes, we studied the effect of *MXI1* interference on hypoxia induced gene repression (supplementary figure 4). Although silencing of *MXI1* attenuated the repression of 245 out of 409 genes found significantly repressed in control HUVEC cells (supplementary figure 3), the effect was of very small magnitude (mean of differences= +0.06, 95% CI [+0.02,+0.09]). Accordingly, the number and identity of genes significantly repressed by hypoxia remained unaltered in silenced cells. This result is in agreement with other reports (71) and suggests functional redundancy with other transcriptional repressors in the *MAD* family, such as *MXD4*, that are also induced by hypoxia in HUVEC cells. Interestingly, the SIN3A complex also binds to the repressor members of the E2F family (61, 62) including E2F4 and E2F6 that were also found enriched in downregulated genes (figure 3). *E2F4* and *E2F6* mRNAs were not induced by hypoxia in HUVEC (supplementary table 1), but both of them have been implicated in the repression of BRCA1 and RAD51 in hypoxia (72, 73). In addition, a recent report indicates that HIF1A physically interacts with E2F7 to form an active repressor complex that prevents transcription (74). Thus, it is feasible that a similar mechanism is operative in HUVEC cells whereby HIF may directly interact with E2F4/6 and regulate the activity of SIN3A on repressed genes. Future work will address this hypothesis and the role of Mad and E2F family members on hypoxia-induced gene repression. Besides MXI1 and E2Fs we found other factors that, although enriched in downregulated genes, did not pass the stringent threshold of adjusted p-value<0.005. Among them is *bHLHe40* (adjusted p-value=0.0107, Fisher’s exact test for the association with downregulated genes). As discussed before, *bHLHe40* has been shown to mediate the repression of specific genes downstream of HIF. Finally, Repressor Element 1-Silencing Transcription factor (REST) has also been recently proposed to repress several genes in hypoxia (75). Interestingly, REST interacts with SIN3A to repress target genes (76). However, although REST is present in the ENCODE dataset, we did not find enrichment for it in HUVEC cells, suggesting that different DNA-binding repressors might target the SIN3A complex to distinct target genes in a cell-specific manner.

Regardless the specific mechanism, our results demonstrate that SIN3A is required for a complete transcriptional response to hypoxia. *SIN3A* interference not only affected individual genes but also altered paradigmatic hypoxia-regulated processes, such as suppression of oxidative phosphorylation and fatty acid oxidation (figure 3). Interestingly, recent reports in *Saccharomyces* and *Drosophila* (77) demonstrate that loss of SIN3 affect mitochondrial activity, suggesting an evolutionarily conserved role for SIN3 in the control of cellular energy production. Further studies will be required to test the role of SIN3A in the changes induced by hypoxia on mitochondrial function.

Altogether, our results delineate an unanticipated role for the SIN3A corepressor complex in the transcriptional response to hypoxia, highlighting that the regulation of gene expression by environmental oxygen is substantially more complex than has previously been considered. More generally, our findings strongly support a prominent role for SIN3A complex in the up- and downregulation of transcription in response to a wide array of environmental stresses.

## ACCESSION NUMBERS

All data sets generated as part of this study are available at NCBI’s Gene Expression Omnibus (GEO) (43, 44) and are accessible through the following GEO Series accession numbers:

RNA-Seq 4SU-labeled HUVEC three biological replicates three RNA fractions: GSE89831

RNA-Seq HUVEC shRNA Sin3A two biological replicate total RNA fractions: GSE89840

RNA-Seq 4SU-labeled HUVEC shRNA EPAS1 one biological replicate two RNA fractions: GSE89838

RNA-Seq HUVEC shRNA EPAS1/HIF1A/MXI1 one biological replicate total RNA fractions: GSE89839

ChIP-Seq HUVEC EPAS1/HIF1A/ARNT one biological replicate: GSE89836

ChIP-Seq HUVEC SIN3A one biological replicate: GSE103245

The reference Series for all the experiments in this publication is GSE89841

## SUPPLEMENTARY DATA

Supplementary Data are available at NAR online.

## ACKNOWLEDGEMENT

We thank Diego Villar for critical review of the manuscript and suggestions. We also like to thank Ramon Diaz-Uriarte for his advice on statistical analysis and for lending us computing power to perform the analysis of high-throughput data.

